# The Evolution of Covert Signaling

**DOI:** 10.1101/132407

**Authors:** Paul E. Smaldino, Thomas J. Flamson, Richard Mcelreath

## Abstract

Human sociality depends upon the benefits of mutual aid and extensive communication. However, diverse norms and preferences complicate mutual aid, and ambiguity in meaning hinders communication. Here we demonstrate that these two problems can work together to enhance cooperation through the strategic use of deliberately ambiguous signals: covert signaling. Covert signaling is the transmission of information that is accurately received by its intended audience but obscured when perceived by others. Such signals may allow coordination and enhanced cooperation while also avoiding the alienation or hostile reactions of individuals with different preferences. Although the empirical literature has identified potential mechanisms of covert signaling, such as encryption in humor, there is to date no formal theory of its dynamics. We introduce a novel mathematical model to assess when a covert signaling strategy will evolve, as well as how receiver attitudes coevolve with covert signals. Covert signaling plausibly serves an important function in facilitating within-group cooperative assortment by allowing individuals to pair up with similar group members when possible and to get along with dissimilar ones when necessary. This mechanism has broad implications for theories of signaling and cooperation, humor, social identity, political psychology, and the evolution of human cultural complexity.

## INTRODUCTION

Much of the research on human cooperation has focused on the free-rider problem: how to maintain cooperation when individuals’ interests are opposed to those of the group. However, individual interests are often *aligned* with those of the group, and these mutualistic scenarios may be equally important in understanding human social evolution (Skyrms, 2004; Calcott, 2008; Tomasello et al., 2012; Smaldino, 2014). Even without incentives for individuals to defect, mutualism provides a different dilemma. When individuals differ in preferences or norms, it is harder to efficiently coordinate. The formation of reliable expectations of partner behavior that make coordination possible is therefore essential for the evolution of mutualism (Schelling, 1960).

Consider, for example, a couple planning their Saturday. Chris wants to go to the opera; Pat wants to go to the monster truck rally. Each would rather do something with their partner than go it alone, but each has a different preference (Luce & Raiffa, 1957). If such mismatches are sufficiently frequent, Chris and Pat might be better off finding new partners with better-aligned interests. Successful cooperation requires resolution of this clash of preferences. Human societies are replete with dilemmas of this kind (Boyd & Richerson, 1994), and the need to efficiently coordinate extends to many forms of collective action (Ostrom, 2000). Institutions like punishment convert other social dilemmas into coordination dilemmas, expanding their importance. If individuals could assort on preferences and norms, cooperative payoffs may be increased. But often these traits are impossible to directly observe. When preferences are consciously held, individuals can merely signal them. But often individuals are not conscious of their preferences or realize their relevance too late to signal them.

One solution is the evolution of ethnic marking. Anthropologists have long argued that ethnic markers may signal group membership and improve cooperative outcomes (Barth, 1969). An extensive formal literature has developed exploring how arbitrary signals can facilitate assortment on unconscious norms and preferences (Castro & Toro, 2007; Efferson et al., 2008; Mace & Holden, 2005; McElreath et al., 2003; Moffett, 2013; Nettle & Dunbar, 1997). Language can also serve as a marker for social coordination (Nettle & Dunbar, 1997), as can visible purchasing and fashion choices (Berger & Heath, 2008; Smaldino et al., 2017).

Communication is implicated in all these solutions. However, much communication is ambiguous. Is this ambiguity merely the result of constraints on the accuracy of communication? It may naïvely appear that communication should have clarity as its goal. However, purposeful ambiguity may allow signalers flexibility and plausible deniability (Eisenberg, 1984; Pinker et al., 2008; Santana, 2014). Previous work has illustrated how leaders may use ambiguous language to rally diverse followers (Eisenberg, 1984), politicians may use vague platforms to avoid committing to specific policies (Aragonès & Neeman, 2000), and suitors may mask their flirtations to be viewed innocuously (or at least to provide plausible deniability) if their affections are unreciprocated (Pinker et al., 2008; Gersick & Kurzban, 2014).

We propose more broadly that ambiguity may enable coordination and thereby enhance cooperation. Consider again the use of signals for assortment on cooperative norms. Overt signals like ethnic markers are useful in some contexts, where the adaptive problem is to delimit a set of partners who subscribe to the same broad behavioral norms and to categorically avoid interaction with those who do not. However, the “all-or-nothing” character of such signals can also foreclose valuable partnerships in different contexts. Signals that communicate similarity can also communicate difference, and this can be damaging for within-group cooperation. Individuals may benefit from not foreclosing relationships with less similar group members, so as to successfully cooperate with them in other contexts. Although any two individuals within a group can cooperate when it is mutually beneficial, pairs who are more similar can cooperate more effectively, generating larger benefits (Kaufman, 1967; Wolosin, 1975; Fischer, 2009; Hruschka, 2010; Toma et al., 2012). To be clear, we focus on those aspects of individual variation for which similarity enhances cooperation—these include norms, values, and identity (Smaldino, 2018a,b). Similarity on these dimensions matter even when the cooperative benefit is maximal for diverse rather than homogenous groups (Hong & Page, 2004). Scenarios in which individuals are unable to effectively assort on norms or attitudes are common, especially in complex societies (for example in business or education settings), but also in smaller societies. A signaling system that enables group members to communicate relative similarity only when similarity is high while retaining a shroud of ambiguity when similarity is low is likely to have been advantageous for much of human history.

*Covert signaling* is the transmission of information that is accurately received by its intended audience but obscured when perceived by others. A common example is “dog-whistling,” in which statements have one meaning for the public at large and a more specialized meaning for others (López, 2014). Such language attempts to transmit a coded message while alienating the fewest listeners possible. A possibly much more common form of covert signaling is humor. According to the encryption model of humor (Flamson & Barrett, 2008; Flamson & Bryant, 2013), a necessary component of humorous production is the presence of multiple, divergent understandings of speaker meaning, some of which are dependent on access to implicit information. Only listeners who share access to this information can “decrypt” the implicit understandings and understand the joke. Because the successful production of a joke requires access to that implicit information, humor behaves in manner similar to “digital signatures” in computer cryptography, verifying the speaker’s access to that information without explicitly stating it. Laughter may also serve as an honest signal of a shared sense of humor (Bryant & Aktipis, 2014), and thus of similarity on a variety of traits. While not all humor has this form, a substantial amount of spontaneous, natural humor does (Flamson & Barrett, 2008; Flamson & Bryant, 2013). We allow that other types of identity signals (*sensu* Berger & Heath 2008) may be also be covert.

We propose that covert signaling serves an important function in facilitating effective cooperation within groups by allowing individuals to assort on norms when possible while avoiding conflict with dissimilar individuals when their assistance is necessary. In the remainder of this paper, we define the logic of covert signaling. We analyze the conditions for selection to favor covert signaling over overt signaling, in which information about an individual’s traits is more transparent. Using a formal model, we show that covert signaling can be favored. It sacrifices transparency for the sake of maintaining working relationships with dissimilar individuals. Although covert signals are less accurate than overt signals, we show that the increased ambiguity can in some cases be advantageous. This characterization of covert signaling in terms of cooperative assortment may therefore help to explain forms of communication and coordination such as coded speech and humor, as well as for the flexibility of human sociality more generally. Nevertheless, covert signaling is not always advantageous. For example, if it is possible to freely choose cooperative partners from a large pool, overt signaling may be more advantageous, as individuals will be better able avoid dissimilar partners. Our model yields specific predictions about default attitudes toward strangers in the absence of clear signals, with implications for understanding differences between contemporary political affiliations.

## MODEL DESCRIPTION

We consider a large population of individuals who have already solved the firstorder cooperation problem of suppressing free riders, and can instead focus on maximizing the benefit generated by cooperation. Although individuals all belong to the same group, they also vary along many trait dimensions and thus share more in common with some individuals than others. Pairs of individuals whose trait profiles overlap to some threshold degree are deemed *similar*. Otherwise they are deemed *dissimilar*. Pairs of similar individuals can more effectively coordinate and obtain higher payoffs. The probability that two randomly selected individuals have similar trait profiles is given by *s*.

Our model proceeds in discrete generations, with each generation subdivided into two stages. In the first stage, individuals signal information about their trait profiles to the other members of the group, and those who receive signals form attitudes about their senders. In the second stage, individuals interact in one of two ways and receive payoffs conditional upon similarity as well as attitudes formed in the first stage.

**Stage 1: Signaling.** Individuals produce either an overt or covert signal of their underlying traits. We study a family of continuous signaling strategies in which covert signals are produced a fraction *p* of the time. Overt signals are received by a fraction *R* of the population and explicitly signal similarity or dissimilarity. Covert signals in contrast are received by a fraction *r* < *R* of the population and have content contingent upon the similarity of the sender and receiver. See Figure 1. When the sender and receiver are similar, covert signals are received as signaling similarity. Otherwise, the receiver does not notice the signal and acts as if no signal was received at all.

**Figure 1.**
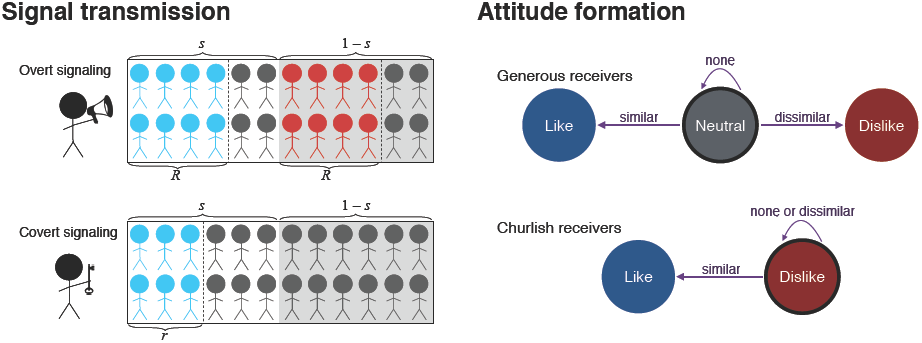
Illustration of the signal transmission and attitude formation components of the model. Left: A proportion *s* of the population is similar. Overt signals are received by a proportion *R* of the total population, while covert signals are received by a proportion *r* < *R* of similar individuals only. Right: Generous receivers default to a neutral attitude in the absence of a signal, while churlish receivers dislike the signaler unless they know he or she is similar.

Receivers have a default attitude towards all individuals in the population and update this attitude upon receiving a signal. Receiver strategy maps three signal states—similar, dissimilar, no signal—to an attitude. We consider three discrete attitudes: like, dislike, and neutral. These correspond to the hypothesis that covert signals help individuals to avoid being disliked while also achieving sufficient positive assortment by type. Two attitudes would be too few, because it would force agents to adopt either like or dislike as a default attitude, removing any incentive for covert signals. Three is the minimum required to model the hypothesis. We allow a continuous family of receiver strategies. Each strategy parameter *a*_xy_ indicates the probability of mapping signal X∈ {Similar,None,Dissimilar} to attitude Y ∈ {Like,Neutral,Dislike}. The total receiver strategy can be represented by a table:

|  | Like | Neutral | Dislike |
| --- | --- | --- | --- |
| Similar | $a_{SL}$ | $a_{SN}$ | $a_{SD}$ |
| None | $a_{NL}$ | $a_{NN}$ | $a_{ND}$ |
| Dissimilar | $a_{DL}$ | $a_{DN}$ | $a_{DD}$ |

The three parameters in each row are constrained to sum to one. Although receiver strategies can vary continuously, we can think of the space of possible strategies as bounded by two extremes. *Generous* receivers default to a neutral attitude in the absence of a signal, while *churlish* receivers dislike the signaler unless they receive a signal of similarity. As we will see, these default receiver strategies are very important.

**Stage 2: Interaction.** After attitudes are established, pairs of individuals interact. There are two interaction contexts, each with its own mode of dyad formation. In a *free choice* scenario, dyads form conditional on the attitudes of both individuals. In contrast, in a *forced choice* scenario, an individual must seek help from whomever happens to be around and dyads are not conditional upon shared attitudes. Under these circumstances, it may be important not to have burned bridges, since this will limit the likelihood of effective coordination.

In the free choice context, dyads form based on joint attitudes, but attitudes do not directly influence payoffs. Instead, payoffs are determined by the underlying similarity of the dyad. Specifically, similar dyads receive an average payoff of 1, establishing a baseline measurement scale. Dissimilar dyads receive a payoff of zero. An individual in a dyad who *likes* the other individual increases the proportional odds of that dyad forming by a factor *w*_L_ > 1. For each individual who *dislikes* the other, the proportional odds of the dyad forming are reduced by a factor *w*_D_ < 1. This implies five possible kinds of dyads that might interact: LL, LN, NN, ND, and DD. The proportional odds of each, relative to random assortment, are: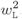, *w*_L_, 1, *w*_D_, and 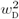 These parameters are fixed features of the social environment, not aspects of strategy. This prevents strategy dynamics from generating perfect assortment. Receiver strategies that assign attitudes in ways that make good use of signal information will achieve better assortment and receive higher payoffs, conditional on the assortment constraints determined by *w*_L_ and *w*_D_.

In the forced choice context, dyads form at random with respect to attitudes, but attitudes directly influence payoffs. This context entails a baseline payoff of 1 for both individuals. However, attitudes adjust payoffs, because negative attitudes make it harder to interact. When one individual dislikes the other, he makes the interaction more difficult than it must be and thereby imposes a cost –*d* on the other individual. When both individuals dislike one another, their difficulties act synergistically, inducing an additional cost –*δ* on each. This cost could result from spite or from uncontrollable inefficiency, a negative consequence of second-order common knowledge (*sensu* Chwe 2001). As we show later, these synergistic costs are very important to the overall signaling dynamics.

Let *q* be the relative importance of the free choice context and 1– *q* the relative importance of the forced choice context. These two contexts are starkly different, presenting the clearest investigation of the hypothesis that covert signals trade worse performance in assortment contexts, in which norms influence payoffs, for better performance in forced contexts in which attitudes influence payoffs. Real contexts are some mix of these extremes, and the parameter *q* allows us to explore the range of mixes.

**Payoff expression.** With the assumptions above, we can define a general payoff expression for a rare individual with signal strategy *p′* and receiver strategy matrix **a***′* in a population with common-type strategy {*p*, **a}**. The expected payoff to this individual is:

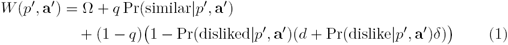

where Ω is an expected payoff due to other activities. The work lies in defining the probabilities Pr(similar *p′*, **a***′*), Pr(dislike *p′*, **a***′*), and Pr(disliked *p′*, **a***′*). In the mathematical appendix, we show how to define these probabilities, using the assumptions above. The resulting general payoff expression is very complicated. In the following section, however, we are nevertheless able to analyze it by considering invasion and stability of relevant combinations of signaling and receiver strategies.

## ANALYSIS AND RESULTS

The motivating hypothesis is that covert signals can proliferate because they allow sufficient assortment in the free choice context and also reduce being disliked in the forced choice context. To evaluate the logic of this idea, we proceed by asking when covert signals can be stable, when they can invade, and which receiver strategies are necessary for their stability or invasion. The following conditions favor covert signals.

1. Covert signals require a sufficient proportion of receivers to default to neutral attitudes. If everyone defaults to disliking, then covert signals can produce no benefit. Defaulting to neutral is favored under a wide range of conditions, provided that covert signals are sufficiently hard to receive (*r* is not too large) and avoidance of disliked individuals is not too efficient (*w*_D_ is not too small).
2. Covert signals require that the cost of being disliked in the forced choice context be sufficiently high. This also means that baseline similarity in the population (*s*) must be sufficiently low, because this creates the risk of being disliked by dissimilar individuals.
3. Overt signalers cannot have too large an advantage in the free choice context. This requires that assortment with liked individuals not be too accurate. The accuracy of assortment is influenced by the reception probabilities of both signal types, *R* and *r*, as well as the proportional odds assortment factors, *w*_L_ and *w*_D_.

In the remainder of this section, we derive these results and provide intuition for why they hold. First we derive simple evolutionary dynamics for these payoffs. This allows us to submit the model to invasion and stability analysis, asking both when covert signals can be stable and when they may invade a population of overt signals. Then we proceed by considering the dynamics within each interaction context—the forced choice context and the free choice context—separately. Then we summarize the joint dynamics of the full model with both contexts.

**Evolutionary dynamics.** We generate evolutionary dynamics for the strategy space by assuming that rare invading strategies increase in frequency when they achieve higher payoffs than a common-type strategy. We define selection gradients for both *p* and the attitude parameters. The gradient for signaling is defined by:

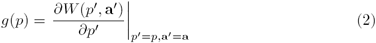

The gradient for each receiver parameter is defined similarly. A number of different mechanisms can generate such dynamics. For example, an individual could acquire its strategy from successful individuals through learning. Genetically coded strategies that influence biological fitness would also generate this dynamic. We remain agnostic about inheritance and transmission mechanism, because the point of our modeling exercise is to explore the design aspects of covert signals. This is best achieved by a form of analysis that abstracts away from transmission details, even though of course in any real system such details will turn out to influence which strategies are possible and how they evolve (Grafen, 1984). We also note that if the mechanism of transmission is cultural, replicators are not strictly necessary but can serve as meaningful approximations of lower-fidelity transmission channels (Henrich & Boyd, 2002).

The potential space of receiver strategies is very large. However, the *relevant* space of strategies is fairly small. In the mathematical appendix, we show that payoff dynamics always favor mapping *similar* signals to *like* attitudes, implying *a*_SL_ = 1. The reason is that maximizing the probability of assortment for similar individuals also maximizes payoffs, and the *like* attitude maximizes the probability of interacting in the free choice context. On the other hand, payoff dynamics do not always favor mapping *dissimilar* signals to *dislike* attitudes. The reason is that the forced choice context disfavors disliking whenever *δ* > 0. We therefore constrain further analysis to the relevant situations in which the penalty for mutual dislike, *δ*, is small enough that assortment incentives favor mapping *dissimilar* signals to *dislike* attitudes. We reemphasize this constraint in the discussion, because constraints of this sort help in producing predictions. Finally, with respect to default attitudes, formed when no signal is received, payoff dynamics never favor assigning *like*, because this erodes the value of assigning *like* to *similar* signals.

The remaining default receiver parameters are free to evolve. Therefore, for most of the analysis to follow, we assume that *a*_SL_ = 1, *a*_DD_ = 1, *a*_NL_ = 0, *a*_NN_ = 1 – *α*, and *a*_ND_ = *α*. This allows us to use the gradient on *α*, defined by:

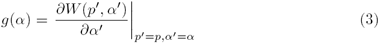

to ask when evolution favors assigning *dislike* to no signal, *α* > 0, effectively defaulting to disliking everyone. As noted earlier, it will be convenient to refer to *α* = 0 as the *generous receiver* strategy and *α* = 1 as the *churlish receiver* strategy.

**Dynamics of the forced choice context.** In this context, incentives favor covert signals, because such signals are better at avoiding being disliked. However, this advantage depends upon incentives favoring generous receiver strategies that do not dislike by default. That said, receiver incentives in this context *always* favor generous receiver strategies as long as there is any negative synergy, *δ* > 0. Therefore the forced choice context favors generous receivers, *α* = 0, which in turn favor covert signals, *p* = 1.

The gradients in this context are:

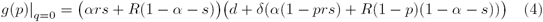

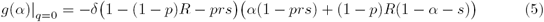

These expressions seem complex at first, but produce fairly simple dynamics. First let’s ask when *p* can increase. When *α* = 0, the generous receiver strategy is common, and covert signals can increase when:

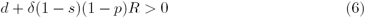

This is satisfied for any allowable values of the parameters. Note also that it does not require both a direct cost of being disliked, *d*, and a synergistic cost of mutual dislike, *δ*. Either one is sufficient to favor covert signals, as long as *α* is small. Next consider when *α* = 1, the churlish receiver strategy is common. Then covert signals can increase when:

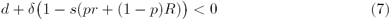

And this is never satisfied, for any *p*. Therefore covert signals are favored when 1-*α*, the amount of generous receiver behavior, is sufficiently high. The threshold value is found where *g*(*p*) = 0:

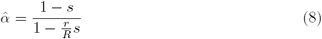

When *α* is above this value, overt signals are favored. When it is below it, covert signals are favored. Why? When receivers are relatively generous, and there is sufficient dissimilarity in the population, covert signals reduce costs by avoiding being disliked. If generous receivers are relatively rare, however, then covert signals can actually do worse than overt signals, because they are received less often than overt signals, *r* < *R*. If *r/R* is sufficiently small, overt signals are favored for a wide range of values of *α*. If however *r* = *R* and covert signals have no disadvantage in audience size, then overt signals are never favored in this context, no matter the amount of similarity *s*.

Now the crucial question is when *α* will fall below this threshold 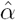. The condition for payoff dynamics to favor smaller values of *α*, in the forced choice context only, is just *δ* > 0. Therefore the forced choice context always favors smaller values of *α*, provided there is any negative synergy between disliking and being disliked. Otherwise *α* is neutral and does not move at all, based on payoff dynamics. Why does this context always favor generous receivers? There is no advantage to be had in disliking people in this context, because assortment does not depend upon attitudes. Payoffs depend upon attitudes, however, and mutual dislike results in poor payoffs. Therefore, it pays to be generous in attitudes towards those one has no information about.

**Dynamics of the free choice context.** In the free choice context, attitudes influence assortment but do not directly influence payoffs. Instead, hidden norm similarity influences payoffs. The free choice context favors overt signals over covert signals, because overt signals increase assortment—such signals are easier to receive and more effectively discriminate similarity. The churlish receiver strategy, *α* = 1, is favored by this context, because it also increases assortment with similar individuals. Therefore this context is hostile to covert signals and to the receiver strategy that favors them.

To support the above statements, we demonstrate that the gradient for *p* in this context is always negative and that the gradient for *α* in this context is always positive. The gradients in this context are:

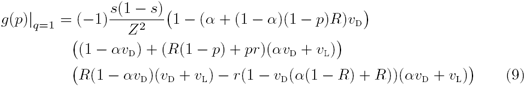

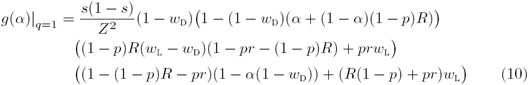

where *Z* is a normalizing term and the symbols *v*_D_ = 1 *w*_D_ and *v*_L_ = *w*_L_ - 1 are used for compactness of notation. By inspection, every term after the leading (–1) in *g*(*p*) is positive, for all allowed values of the variables, and so the gradient is always negative. Similarly, every term in *g*(*α*) is positive, and so the gradient is always positive. Therefore payoff incentives in the free choice context never favor covert signals and always favor churlish receivers.

While this context always favors overt signalers and churlish receivers, the strength of the incentives may vary. First, both gradients are proportional to the variance in similarity, *s*(1 – *s*). This indicates that intermediate similarity more strongly favors overt signals, unlike the situation in the forced choice context, in which high similarity favored overt signals. The reason that the variance matters now is that payoffs depend directly upon similarity, not upon attitudes. The more variance in similarity in the population, the greater the advantage of efficient assortment.

Overt signals have the advantage in this context, because they are better at assortment. Therefore, any change in variables that reduces the accuracy of assortment overall will reduce overt signalers’ advantage. The important variables are *R*, the probability an overt signal is received, and the assortment proportional odds *w*_D_ and *w*_L_. Reducing *R* reduces overt signalers’ advantage, because it makes signals less valuable overall. Making either *w*_D_ or *w*_L_ closer to 1 makes assortment, based on attitudes, less effective. This also reduces overt signalers’ advantage.

**Joint dynamics: When do covert signals evolve?** When both contexts matter, the joint dynamics take on one of three characteristic regimes. First, covert signals both invade and are evolutionarily stable. Second, overt signals both invade and are evolutionary stable. Third, a mixed equilibrium exists at which covert and overt signals coexist in the population. Figure 2 illustrates these three regimes. These three examples all weigh the forced choice and free choice contexts equally, *q* = 0.5. The other parameters then shift the strength of incentives in each context to influence overall dynamics. For other values of *q*, the strength of incentives would have to shift as well to overcome weight given to each context.

**Figure 2.**
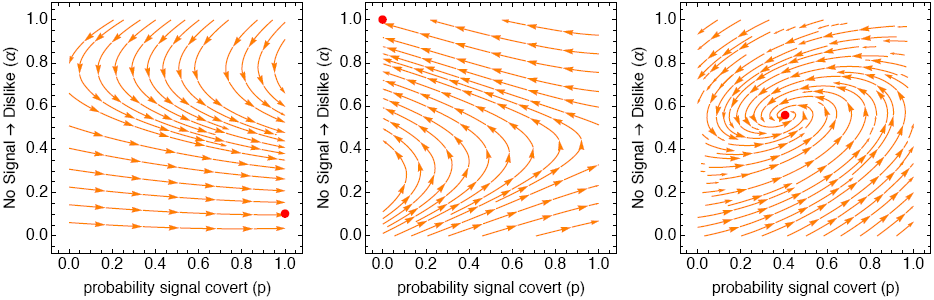
The three dynamic regimes that arise from the joint dynamics. In each plot, the paths show the evolutionary trajectories in each region of the phase space defined by the probability of covert signals (*p*, horizontal) and the probability of churlish receivers (*α*, vertical). The red points show equilibria. In all three plots: *d* = 0.1, *δ* = 0.01, *q* = 0.5. Left: *s* = 0.1, *r/R* = 0.25, *w*_L_ = 1.1, *w*_D_ = 0.9. Middle: Same as left, but *w*_D_ = 0.6. Right: *s* = 0.2, *r/R* = 0.75, *w*_L_ = 1.25, *w*_D_ = 0.8.

When covert signals are sufficiently noisy (*r/R* low), similarity is sufficiently rare (*s* low), and assortment (*w*_L_, *w*_D_) not too efficient, covert signals can both invade and are an ESS. This situation is shown in the lefthand plot. While dynamics do not favor covert signals when *α* is large, near the top of the phase space, dynamics in that region favor smaller values of *α*. Eventually, *α* becomes small enough to allow covert signals to invade and reach fixation. In many cases, a small amount of churlish receiver strategy, *α >* 0, persists.

When the conditions outlined above are not met, incentives favor instead overt signals. The middle plot illustrates a case essentially the opposite of the one on the left. Here, *w*_D_ = 0.6, making assortment efficient. When assortment is efficient, it may pay to dislike by default. This sets up a dynamic that eventually favors overt signals. While covert signals are still favored when *α* is low, the fact that larger values of *α* are favored everywhere leads eventually to invasion and fixation of overt signals.

Finally, the plot on the right shows an intermediate case, in which conditions favor both signaling strategies. Here *s* = 0.2, *r/R* = 0.75, *w*_L_ = 1.25, and *w*_D_ = 0.8. In this regime, the conditions that favor covert signals also favor more churlish receivers. Similarly, the conditions that favor overt signals also favor fewer churlish receivers. In total, the population comes to rest with a mixture of signaling and receiving strategies.

A more general view of the dynamics is available by considering the boundary conditions that make covert signaling an ESS. Recall that *q* is the relative importance of free choice scenarios. Define a threshold 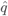 as the largest value of *q* for which covert signals can resist invasion by overt signals. This is defined by values of 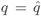 and 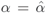 that satisfy *g*(*p*) |*p=*1 = 0 and *g*(*α*) |*p=*1 = 0. These cannot in general be solved analytically. So we solve the system numerically, in order to illustrate the range of joint dynamics. Values of *q* less than 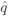 make covert signals evolutionarily stable—overt signals cannot invade. Values of *q* greater than 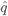 allow overt signals to invade, though we note that covert signals may nevertheless remain in the population, at an internal stable value *p*. Therefore 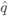 provides a useful metric of how strongly a parameter configuration favors covert signals.

We use 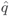 to summarize the tradeoffs in the signaling model. Recall that the cost of being disliked, *d*, is needed to favor covert signals. Therefore increasing *d* makes it easier for covert signals to be an ESS. However, the rate of similarity, *s*, favors overt signals. It is of value to note that *d* cannot compensate for *s*—if *s* is large, then steeply increasing costs *d* will not favor covert signals. We show this relationship in Figure 3, lefthand plot. Each curve in this plot is a threshold 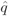, below which covert signaling is an ESS. For each level of *s*, the impact of increasing *d* diminishes rapidly. Therefore some cost *d* is necessary for covert signals to evolve and be stable, but these costs cannot easily compensate when similarity is sufficiently common.

**Figure 3.**
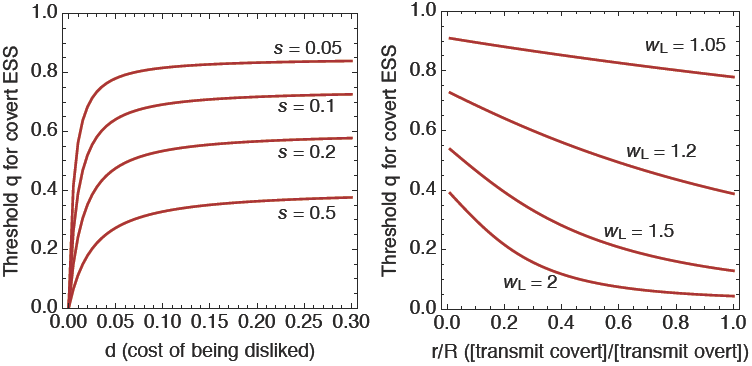
Plots of the largest value of *q*, 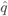, that allows covert signaling to be an ESS. Each curve represents a set of parameter values. Points below each curve make covert signals uninvadvable by overt signals. Points above each curve allow overt signaling to invade. Left: The cost of being disliked, *d*, for four values of the baseline rate of similarity, *s*. *r/R* = 0.5, *w*_L_ = 1.1, *w*_D_ = 0.9, and *δ* = 0.01. Right: The ratio of covert transmission rate, *r*, to the rate of overt transmission rate, *R*, for four values of *w*_L_ = 1*/w*_D_. *s* = 0.1, *R* = 0.5, *d* = 0.1, and *δ* = 0.01.

Consider another important pair of dimensions: the ratio of transmission rates *r/R* in covert/overt signals and the efficiency of assortment, as measured by *w*_L_ and *w*_D_. Figure 3, righthand side, shows 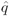 curves for four values of *w*_L_ = 1*/w*_D_, as functions of *r/R*. Covert signals are favored when *r/R* is small, as explained in the previous sections. But when assortment is very efficient, such as *w*_L_ = 2 near the bottom of the plot, it requires very low values of *r/R* to compensate in favor of overt signaling.

## DISCUSSION

The dynamics of cooperation are more complicated than implied by models in which maximal benefits accrue to those who can simply avoid free riders. Not all cooperators are equal. Individuals vary, making assortment among cooperators important. Circumstances also vary. When individuals must occasionally collaborate with those outside their circles of friends, it can be critical to avoid burning bridges with dissimilar members of one’s group. Covert signaling makes this possible, and this may be why phenomena like humor are observed in all human societies, at both small and large scales (Apte, 1985; Brown, 1991).

We have shown that covert signaling is favored when forced choice scenarios are common, when similarity is low, when the cost of being disliked is high, and when covert signals are sufficiently noisy to make the meaning of a signal’s absence ambiguous. We emphasize that covert signaling can be favored even though it is less effective than overt signaling at communicating similarity, because it simultaneously avoids communicating dissimilarity. Although we have focused our attention on the initial establishment of cooperative relationships via signaling, we also note that people can change over time; they may grow more similar to one another or further apart. Covert signals may be important for the continued maintenance of a relationship, or for its reestablishment after prolonged absence.

Our model points to interesting transitions from inter-to intra-group assortment dynamics. As noted, overt signaling systems are favored when the ability to assort on attitudes is high, and when being disliked by dissimilar individuals carries little risk. This is precisely the kind of situation that is assumed to obtain in inter-group assortment, where overt signals such as ethnic markers are used to discriminate between similar and dissimilar individuals. In these between-group contexts, the difference between similar and dissimilar individuals is so great that attempting to coordinate with dissimilar others is not worth the effort, and one can afford to burn bridges with them in order to ensure that similar others are aware of their similarity (McElreath et al., 2003). In fact, it might be argued that burning bridges with dissimilar out-group members is as much a goal of overt signals like ethnic markers as is attracting similar in-group members.

Intra-group assortment, however, is not simply a matter of scaling down inter-group dynamics. Rather, we must already presume some baseline level of similarity resulting from inter-group assortment; for there to be a group within which to assort, some degree of similarity should already be in place that defines that group, such as the shared interaction norms, communication systems, etc. that ethnic markers are thought to ensure. The benefits of further assorting on the basis of more nuanced similarity are therefore likely to be marginal relative to random assortment within the group. When such benefits are small but the costs of being disliked are high, covert signaling is favored.

Relatedly, we emphasize that the probability of similarity, *s*, need not reflect some number of discrete types in the population, but can instead refer to a level of selectivity in how much a given pair of individuals must have in common to be considered “similar.” That is, *s* refers to the proportion of the population that could be considered similar to a focal individual in a given context, with higher values indicating a looser concept of “similar” than lower ones. Changes to *s* can have a significant impact on the overall dynamics of the system. When *s* is large, the focus is on avoiding rare dissimilar individuals, and overt signals will be favored. As *s* decreases, the criteria for considering a potential partner sufficiently similar to reap the benefits of enhanced coordination become stricter, and the utility of covert signals increases.

An interesting direction for future exploration is how these dynamics might respond to increased social complexity. In the larger and more complex societies associated with the development of agriculture, and particularly in the last few centuries, interactions with strangers are more frequent and occur across many contexts, necessitating strategies for temporary assortment (Johnson & Earle, 2000; Smaldino, 2018b,a). Consequently, expected similarity will be lower, signal fidelity will be noisier, and assortment on attitudes will be less efficient. These are precisely the conditions in our model associated with the evolution of covert signaling. In large, diverse populations, covert signaling may sustain social cohesion and prevent burning bridges between individuals or groups that must occasionally collaborate. That said, covert signals are not necessarily rare in small-scale societies. Our own experiences in the field and conversations with other researchers indicate that they occur with some regularity. Our model can help to identify contexts in which covert signaling should or should not be expected.

Identity signaling, whether overt social markers or more covert communication, can be used by individuals looking to find others similar to themselves and to avoid being mistaken for something they are not (Berger & Heath, 2008; Smaldino, 2018b,a). If the need to cooperate with dissimilar individuals is unlikely or if similar individuals are common, then overt declarations of identity should be expected. On the other hand, if burning bridges is both costly and likely given an overt signaling strategy, we should expect identity to be signaled much more subtly. In reality, increasing levels of specificity may be signaled in increasingly covert ways, and without all received signals actively inducing a change in disposition toward the sender. A related signaling strategy, not covered by our model, might facilitate liking between similar individuals but only indifference otherwise. Casual, coarsegrain identity signaling may often take this form, as in cases of fashion adoption or pop culture allegiances. It would be interesting to investigate how common these “semi-covert” signals are in small-scale communities, as they seem pervasive in complex industrialized societies.

Our model additionally helps make sense of findings from political psychology suggesting that people in the industrialized West who identify as conservative or right-leaning tend to view ambiguous people as hostile, while those identifying as liberal or left-leaning tend view ambiguous people as neutral (Vigil, 2010; Hibbing et al., 2014; Holbrook et al., 2017). In our model, a default attitude to dislike was linked with overt signaling, which we in turn associate with the preservation of strong between-group boundaries. In contrast, a default attitude of neutral was associated with covert signaling, and with the avoidance of burned bridges to facilitate more widespread within-group cooperation. As a broad generalization, our analysis suggests that conservatives may be operating under the assumptions of stronger ingroup/outgroup boundaries, increased expectations of similarity toward those they signal, and lower costs to being disliked by dissimilar individuals. In contrast, liberals may be operating under the assumptions of a more broadly defined ingroup, limited expectations for similarity toward those they signal, and higher costs to being disliked by dissimilar individuals. Lending modest support to this idea is the finding that conservatives appear to have a stronger “need for cognitive closure” (reviewed in Hibbing et al., 2014), which is associated with, among other things, a distaste for uncertainty and ambiguity. The modeling framework we present in this paper may thus be useful in understanding patterns of differences between groups, including but not limited to political affiliation.

Ours is the first formal model of covert signaling. As such, it necessarily involves simplifying assumptions concerning the nature of signaling and cooperative assortment. For example, while we have allowed for covert signaling errors in the form of failed transmission to similar individuals, we have not included the converse form of error, where dissimilar individuals *are* able to detect the signal some of the time, and therefore update their disposition to disliking the covert signaler. Adding an additional parameter to account for this possibility does not qualitatively change our analysis. But it may create conditions where a non-signaling “quiet” strategy could invade. In addition, we ignore the possibility of strategic action on the part of the receiver to either improve coordination or to avoid partnering with dissimilar individuals entirely. We assumed that a pairing of dissimilar partners would simply lead to an unsuccessful collaboration, but such a pairing might instead lead each individual to pursue more individualistic interests. At the population level, we assumed that all individuals had an equal probability of encountering similar individuals, and that all similar and dissimilar individuals were equivalent. In reality, some individuals may be more or less likely to encounter similar individuals, perhaps related to differences in the tendency to be conformity- versus distinctiveness-seeking (Smaldino & Epstein, 2015), or reflecting minority-majority dynamics (Wimmer, 2013; Bunce & McElreath, 2018). Exploration of this variation opens the door to evaluating signaling and assortment strategies in stratified groups. All of these limitations provide avenues for future research that will build upon the central findings reported here.

In a population where individuals vary and burning bridges is costly, overtly announcing precisely where one stands entails venturing into a zone of danger. Covert signaling, as in the case of humor or otherwise encrypted language, allows individuals to effectively assort when possible while avoiding burned bridges when the situation calls for partnerships of necessity.

## ACKNOWLEDGMENTS

Thanks to Patrick Barclay, William Baum, John Bunce, and several anonymous reviewers for helpful comments. This work was supported by National Science Foundation Grant BCS 1357240 (to T.F. and R.M.) and by the Division of Social Sciences Dean’s Office at the University of California Davis.

## APPENDIX (ONLINE SUPPLEMENT) The Evolution of Covert Signaling APPENDIX A. MODEL DERIVATION

**A.1 Payoff expressions.** As explained in the main text, we can define a general payoff expression for an individual with strategy *p′* and attitude matrix **a***′* in a population with strategy *{p*, **a***}*. This expression is:

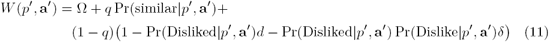

where Ω is an expected baseline payoff due to other activities.

**A.2 Context 1 probabilities.** Pr(similar*|p′*, **a***′*) is the probability of ending up in a similar dyad, given the focal has signaling strategy *{p′*, **a***′}*. By definition:

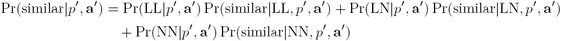

The terms like Pr(similar*|*NN, *p′*, **a***′*) are defined by conditional probaility:

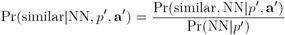

Defining the two terms on the right requires defining probabilities for dyad formation.

The probability that an LL dyad forms is:

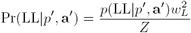

where the denominator *Z* normalizes the probability and *p*(LL *p′*, **a***′*) is the raw proportion of dyads that are LL, post signaling. Under the baseline receiver strategy, **<u>a</u>***′*, it is defined as:

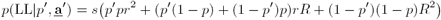

While we later develop this expression in general for all receiver strategies, it’s worth consider the specific expression above, for sake of comprehension. To understand this expression, consider that LL dyads must comprise similar individuals, and both signals have to be received for the attitudes to form. Individuals are similar *s* of the time. There are three ways this can happen: (1) both individuals signal covertly *p′p* of the time, (2) one individual signals covertly and the other overtly *p′*(1 – *p*) + (1 – *p′*)*p* of the time, or (3) both individuals signal overtly (1 – *p′*)(1 – *p*) of the time. In all three cases, both signals must be received. Probabilities for each of the other dyad types—LN, NN, ND, and DD—are defined similarly. Further down, we define all of these probabilities in general, using a more algorithmic approach.

The denominator *Z* normalizes the probabilities of each dyad forming. It is merely the sum of all of the numerators in the probabilities of different types of dyads. These other probabilities are defined similarly:

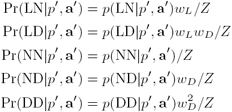

**A.3 Building generalized probabilities.** Now we construct the general probabilities that comprise the payoff expression by using a table of interactions.

| [1]Focal | [2]Other | [3]Sim | [4]Signals | [5]Probability of dyad | [6]FL | [7]FN | [8]FD | [9]OL | [10]ON | [11]OD |
| --- | --- | --- | --- | --- | --- | --- | --- | --- | --- | --- |
| C | C | 1 | 11 | $p'psr^2$ | $a'_{SL}$ | $a'_{SN}$ | $a'_{SD}$ | $a_{SL}$ | $a_{SN}$ | $a_{SD}$ |
| C | C | 1 | 10 | $p'psr(1-r)$ | $a'_{YL}$ | $a'_{YN}$ | $a'_{YD}$ | $a_{SL}$ | $a_{SN}$ | $a_{SD}$ |
| C | C | 1 | 01 | $p'ps(1-r)r$ | $a'_{SL}$ | $a'_{SN}$ | $a'_{SD}$ | $a_{NL}$ | $a_{NN}$ | $a_{ND}$ |
| C | C | 1 | 00 | $p'ps(1-r)^2$ | $a'_{NL}$ | $a'_{NN}$ | $a'_{ND}$ | $a_{NL}$ | $a_{NN}$ | $a_{ND}$ |
| C | C | 0 | 11 | $p'p(1-s)r^2$ | $a_{SL}$ | $a_{SN}$ | $a_{SD}$ | $a_{NL}$ | $a_{NN}$ | $a_{ND}$ |
| C | C | 0 | 10 | $p'p(1-s)r(1-r)$ | $a'_{YL}$ | $a'_{YN}$ | $a'_{YD}$ | $a_{NL}$ | $a_{NN}$ | $a_{ND}$ |
| C | C | 0 | 01 | $p'p(1-s)(1-r)r$ | $a'_{SL}$ | $a'_{SN}$ | $a'_{SD}$ | $a_{NL}$ | $a_{NN}$ | $a_{ND}$ |
| C | C | 0 | 00 | $p'p(1-s)(1-r)^2$ | $a'_{NL}$ | $a'_{NN}$ | $a'_{ND}$ | $a_{NL}$ | $a_{NN}$ | $a_{ND}$ |
| C | O | 1 | 11 | $p'(1-p)srR$ | $a_{SL}$ | $a_{SN}$ | $a_{SD}$ | $a_{SL}$ | $a_{SN}$ | $a_{SD}$ |
| C | O | 1 | 10 | $p'(1-p)sr(1-R)$ | $a'_{YL}$ | $a'_{YN}$ | $a'_{YD}$ | $a_{SL}$ | $a_{SN}$ | $a_{SD}$ |
| C | O | 1 | 01 | $p'(1-p)s(1-r)R$ | $a'_{SL}$ | $a'_{SN}$ | $a'_{SD}$ | $a_{NL}$ | $a_{NN}$ | $a_{ND}$ |
| C | O | 1 | 00 | $p'(1-p)s(1-r)(1-R)$ | $a'_{NL}$ | $a'_{NN}$ | $a'_{ND}$ | $a_{NL}$ | $a_{NN}$ | $a_{ND}$ |
| C | O | 0 | 11 | $p'(1-p)(1-s)rR$ | $a'_{YL}$ | $a'_{YN}$ | $a'_{YD}$ | $a_{NL}$ | $a_{NN}$ | $a_{ND}$ |
| C | O | 0 | 10 | $p'(1-p)(1-s)r(1-R)$ | $a'_{SL}$ | $a'_{SN}$ | $a'_{SD}$ | $a_{NL}$ | $a_{NN}$ | $a_{ND}$ |
| C | O | 0 | 01 | $p'(1-p)(1-s)(1-r)R$ | $a'_{NL}$ | $a'_{NN}$ | $a'_{ND}$ | $a_{NL}$ | $a_{NN}$ | $a_{ND}$ |
| C | O | 0 | 00 | $p'(1-p)(1-s)(1-r)(1-R)$ | $a'_{NL}$ | $a'_{NN}$ | $a'_{ND}$ | $a_{NL}$ | $a_{NN}$ | $a_{ND}$ |
| O | C | 1 | 11 | $(1-p')psRr$ | $a_{SL}$ | $a_{SN}$ | $a_{SD}$ | $a_{SL}$ | $a_{SN}$ | $a_{SD}$ |
| O | C | 1 | 10 | $(1-p')psR(1-r)$ | $a'_{YL}$ | $a'_{YN}$ | $a'_{YD}$ | $a_{SL}$ | $a_{SN}$ | $a_{SD}$ |
| O | C | 1 | 01 | $(1-p')ps(1-R)r$ | $a'_{SL}$ | $a'_{SN}$ | $a'_{SD}$ | $a_{NL}$ | $a_{NN}$ | $a_{ND}$ |
| O | C | 1 | 00 | $(1-p')ps(1-R)(1-r)$ | $a'_{NL}$ | $a'_{NN}$ | $a'_{ND}$ | $a_{NL}$ | $a_{NN}$ | $a_{ND}$ |
| O | C | 0 | 11 | $(1-p')p(1-s)Rr$ | $a'_{YL}$ | $a'_{YN}$ | $a'_{YD}$ | $a_{DL}$ | $a_{DN}$ | $a_{DD}$ |
| O | C | 0 | 10 | $(1-p')p(1-s)R(1-r)$ | $a'_{SL}$ | $a'_{SN}$ | $a'_{SD}$ | $a_{DL}$ | $a_{DN}$ | $a_{DD}$ |
| O | C | 0 | 01 | $(1-p')p(1-s)(1-R)r$ | $a'_{NL}$ | $a'_{NN}$ | $a'_{ND}$ | $a_{NL}$ | $a_{NN}$ | $a_{ND}$ |
| O | C | 0 | 00 | $(1-p')p(1-s)(1-R)(1-r)$ | $a'_{NL}$ | $a'_{NN}$ | $a'_{ND}$ | $a_{NL}$ | $a_{NN}$ | $a_{ND}$ |
| O | O | 1 | 11 | $(1-p')(1-p)sR^2$ | $a_{SL}$ | $a_{SN}$ | $a_{SD}$ | $a_{SL}$ | $a_{SN}$ | $a_{SD}$ |
| O | O | 1 | 10 | $(1-p')(1-p)sR(1-R)$ | $a'_{YL}$ | $a'_{YN}$ | $a'_{YD}$ | $a_{SL}$ | $a_{SN}$ | $a_{SD}$ |
| O | O | 1 | 01 | $(1-p')(1-p)s(1-R)R$ | $a'_{SL}$ | $a'_{SN}$ | $a'_{SD}$ | $a_{NL}$ | $a_{NN}$ | $a_{ND}$ |
| O | O | 1 | 00 | $(1-p')(1-p)s(1-R)^2$ | $a'_{NL}$ | $a'_{NN}$ | $a'_{ND}$ | $a_{NL}$ | $a_{NN}$ | $a_{ND}$ |
| O | O | 0 | 11 | $(1-p')(1-p)(1-s)R^2$ | $a'_{YL}$ | $a'_{YN}$ | $a'_{YD}$ | $a_{DL}$ | $a_{DN}$ | $a_{DD}$ |
| O | O | 0 | 10 | $(1-p')(1-p)(1-s)R(1-R)$ | $a'_{SL}$ | $a'_{SN}$ | $a'_{SD}$ | $a_{DL}$ | $a_{DN}$ | $a_{DD}$ |
| O | O | 0 | 01 | $(1-p')(1-p)(1-s)(1-R)R$ | $a'_{NL}$ | $a'_{NN}$ | $a'_{ND}$ | $a_{NL}$ | $a_{NN}$ | $a_{ND}$ |
| O | O | 0 | 00 | $(1-p')(1-p)(1-s)(1-R)^2$ | $a'_{NL}$ | $a'_{NN}$ | $a'_{ND}$ | $a_{NL}$ | $a_{NN}$ | $a_{ND}$ |

The columns of this table specify:

1. The focal individual’s signaling strategy
2. The other individual’s signaling strategy
3. Whether or not (0/1) the individual’s are similar
4. Whether or not (0/1) focal/other signals are received by the other individual in the pair. 11 indicates that both signals are received. 10 indicates that focal’s signal was received, while other’s signal was not.
5. The probability of this pairing and pair of signal reception events
6. The probability that the focal individual forms attitude L. When a signal is received from a similar individual, this is *a′*_SL_. When no signal is received or a covert signal is received from a dissimilar individual, this is *a′*_nl_. When an overt signal is received from a dissimilar individual, this is *a′*_dl_.
7. The probability that the focal individual forms attitude N
8. The probability that the focal individual forms attitude D
9. The probability that the other individual forms attitude L
10. The probability that the other individual forms attitude N
11. The probability that the other individual forms attitude D

Parameters marked by a prime, such as *p′* and *a′*_SL_, indicate aspects of the focal individual’s strategy, to be contrasted with population values. Again, we refer to the vector of attitude parameters with **a**.

Call this table **M**. To compute probabilities, we multiply specific terms in each row and then sum these products down the rows. For example, the probability that the focal individual and a random individual mutually like one another, *p*(LL*|p′*, **a***′*), is defined by:

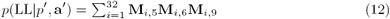

The other probabilities are defined similarity:

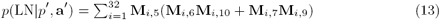

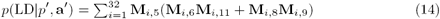

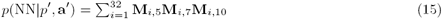

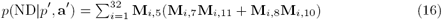

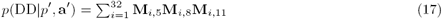

To derive probabilities of similarity and attitudes, all that is required is to multiply each of the products above with the corresponding value in column 3. This defines:

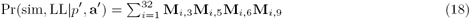

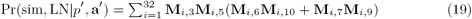

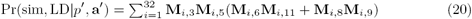

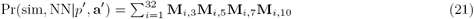

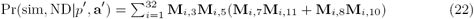

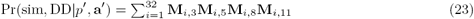

All that remains are probabilities that any random individual Likes or Dislikes the focal:

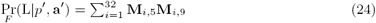

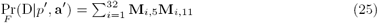

as well as the probabilities that the focal likes or dislikes the other individual:

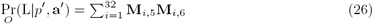

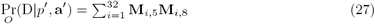

## REFERENCES

Apte, M., (1985) Humor and laughter: An Anthropological Approach. Ithaca, NY: Cornell University Press.

Aragonès, E. & Neeman, Z. (2000). Strategic ambiguity in electoral competition. Journal of Theoretical Politics, 12(2), 183–204.

Barth, F. (1969). Introduction. In F. Barth (Ed.), Ethnic Groups and Boundaries(pp. 9–38). New York: Little, Brown.

Berger, J. & Heath, C. (2008). Who drives divergence? identity signaling, outgroup dissimilarity, and the abandonment of cultural tastes Journal of Personality and Social Psychology, 95(3), 593.

Boyd, R. & Richerson, P. J. (1994). The evolution of norms: nAn anthropological view Journal of Institutional and Theoretical Economics, 150(1), 72–87.

Brown, D. (1991)Human universals. Philadelphia: Temple University Press.

Bryant, G. A. & Aktipis, C. A. (2014). The animal nature of spontaneous human laughter Evolution and Human Behavior, 35, 327–335.

Bunce, J. A. & McElreath, R. (2018). Sustainability of minority culture when inter-ethnic interaction is profitable Nature Human Behaviour.

Calcott, B. (2008). The other cooperation problem: Generating benefit Biology and Philosophy, 23(2), 179–203.

Castro, L. & Toro, M. A. (2007). Mutual benefit cooperation and ethnic cultural diversity Theoretical Population Biology, 71(3), 392–399.

Efferson, C., Lalive, R., & Fehr, E. (2008). The coevolution of cultural groups and ingroup favoritism Science, 321(5897), 1844–1849.

Eisenberg, E. M. (1984). Ambiguity as strategy in organizational communication. Communication Monographs, 51(3), 227–242.

Fischer, I. (2009). Friend or foe: Subjective expected relative similarity as a de-terminant of cooperation Journal of Experimental Psychology: General, 138, 341–350.

Flamson, T. & Barrett, H.(2008). The encryption theory of humor: nA knowledge-based mechanism of honest signaling Journal of Evolutionary Psychology, 6(4), 261–281.

Flamson, T. J. & Bryant, G. A. (2013). Signals of humor: Encryption and laughter in social interaction. In M. Dynel (Ed.), Developments in Linguistic Humour Theory, volume 1 (pp. 49–73). Amsterdam: John Benjamins Publishing.

Gersick, A. & Kurzban, R. (2014). Covert sexual signaling: Human flirtation and implications for other social species Evolutionary Psychology, 12(3), 49–569.

Grafen, A. (1984). Natural selection, kin selection and group selection. In J. Krebs &N. B. Davies (Eds.), Behavioural ecology: An evolutionary approach (pp. 62–84).

Blackwell.Henrich, J. & Boyd, R. (2002). On modeling cognition and culture: Why cultural evolution does not require replication of representations Journal of Cognition and Culture, 2, 87–111.

Hibbing, J. R., Smith, nK. B., & Alford, J. R. (2014). Differences in negativity bias underlie variations in political ideology Behavioral and Brain Sciences, 37, 297–350.

Holbrook, C., López-Rodríguez, L., Fessler, D. M. T., Vázquez, A., & Gómez, A. (2017). Gulliver’s politics: Conservatives envision potential enemies as readily vanquished and physically small Social Psychological and Personality Science, 8(6), 670–678.

Hong, L. & Page, S. E. (2004). Groups of diverse problem solvers can outperform groups of high-ability problem solvers Proceedings of the National Academy of Sciences, 101, 16385–16389.

Hruschka, D. J. (2010)Friendship: Development, ecology, and evolution of a relationship. Berkeley: University of California Press.

Johnson, A. W. & Earle, T. lK. (2000)The Evolution of Human Societies: From Foraging Group to Agrarian State. Stanford University Press.

Kaufman, H. (1967). Similarity and cooperation received as determinants of coop-eration rendered Psychonomic Science, 9, 73–74.

López, I. H. (2014)Dog Whistle Politics: How Coded Racial Appeals Have Reinvented Racism & Wrecked the Middle Class. Oxford University Press.

Luce, R. D. & Raiffa, H. (1957) Games and Decisions.

Wiley.Mace, R. &Holden, C.J. (2005). A phylogenetic approach to cultural evolution Trends in lEcology and Evolution, 20(3), 116–121.

McElreath, R., Boyd, R., & Richerson, P. J. (2003). Shared norms and the evolution of ethnic markers Current Anthropology, 44(1), 122–130.

Moffett, M. W. (2013). Human identity and the evolution of societies Human Nature, 24(3), 219–267.

Nettle, D. & Dunbar, R. (1997). Social markers and the evolution of reciprocal exchange Current Anthropology, 38, 93–99.

Ostrom, E. (2000). Collective action and the evolution of social norms The Journal of Economic Perspectives, 14(3), 137–158.

Pinker, S., Nowak, M. A., & Lee, J. J. (2008). The logic of indirect speechPro-ceedings of the National Academy of Sciences, 105(3), 833–838.

Santana, C. (2014). Ambiguity in cooperative signaling Philosophy of Science, 81,398–422.

Schelling, T. C. (1960) The Strategy of Conflict. Harvard University Press. Skyrms, B. (2004) The Stag Hunt and the Evolution of Social Structure. Cambridge University Press.

Smaldino, P. E. (2014). The cultural evolution of emergent group-level traits Be-havioral and Brain Sciences, 37, 243–295.

Smaldino, P. E. (2018a). The evolution of the social self: Multidimensionality of social identity solves the coordination problems of a society. In A. C. Love & W. C. Wimsatt (Eds.), Beyond the Meme: Development and Structure in Cultural Evolution. University Minnesota Press.

Smaldino, P. E. (2018b). Social identity and cooperation in cultural evolution Behavioural Processes.

Smaldino, P. E. & Epstein, J. M. (2015). Social conformity despite individual preferences for distinctiveness Royal Society Open Science, 2(3), 140437.

Smaldino, P. E., Janssen, M. lA., Hillis, V., & Bednar, J. (2017). Adoption as a social marker: Innovation diffusion with outgroup aversion Journal of Mathematical Sociology, 41, 26–45.

Toma, C., Corneille, O., & Yzerbyt, V. (2012). Holding a mirror up to the self: Egocentric similarity beliefs underlie social projection in cooperation Personality and Social Psychology Bulletin, 38, 1259–1271.

Tomasello, M., Melis, A. P., Tennie, C., Wyman, E., & Herrmann, E. (2012). Two key steps in the evolution of human cooperation Current Anthropology, 53(6), 673–692.

Vigil, J. M. (2010). Political leanings vary with facial expression processing and psychosocial functioning Group Processes & Intergroup Relations, 13(5), 547–558.

Wimmer, A. (2013) Ethnic Boundary Making: Institutions, Power, Networks. Oxford University Press.

Wolosin, R. J. (1975). Cognitive similarity and group laughter Journal of Person-ality and Social Psychology, 32, 503–509.

